# Spike mutation T403R allows bat coronavirus RaTG13 to use human ACE2

**DOI:** 10.1101/2021.05.31.446386

**Authors:** Fabian Zech, Daniel Schniertshauer, Christoph Jung, Alexandra Herrmann, Qinya Xie, Rayhane Nchioua, Caterina Prelli Bozzo, Meta Volcic, Lennart Koepke, Jana Krüger, Sandra Heller, Alexander Kleger, Timo Jacob, Karl-Klaus Conzelmann, Armin Ensser, Konstantin M.J. Sparrer, Frank Kirchhoff

**Affiliations:** Institute of Molecular Virology, Ulm University Medical Center, 89081 Ulm, Germany; Institute of Electrochemistry, Ulm University, 89081 Ulm, Germany; Helmholtz-Institute Ulm (HIU) Electrochemical Energy Storage, Helmholtz-Straße 16, 89081 Ulm, Germany; Karlsruhe Institute of Technology (KIT), P.O. Box 3640, 76021 Karlsruhe, Germany; Institute of Clinical and Molecular Virology, University Hospital Erlangen, Friedrich-Alexander Universität Erlangen-Nürnberg, 91054 Erlangen, Germany; Department of Internal Medicine I, Ulm University Medical Center, 89081 Ulm, Germany; Max von Pettenkofer-Institute of Virology, Medical Faculty, and Gene Center, Ludwig-Maximilians-Universität München, 81377 Munich, Germany

**Keywords:** SARS-CoV-2, RaTG13, spike glycoprotein, ACE2 receptor, viral zoonosis

## Abstract

Severe acute respiratory syndrome coronavirus 2 (SARS-CoV-2), the cause of the COVID-19 pandemic, most likely emerged from bats^1^. A prerequisite for this devastating zoonosis was the ability of the SARS-CoV-2 Spike (S) glycoprotein to use human angiotensin-converting enzyme 2 (ACE2) for viral entry. Although the S protein of the closest related bat virus, RaTG13, shows high similarity to the SARS-CoV-2 S protein it does not efficiently interact with the human ACE2 receptor^2^. Here, we show that a single T403R mutation allows the RaTG13 S to utilize the human ACE2 receptor for infection of human cells and intestinal organoids. Conversely, mutation of R403T in the SARS-CoV-2 S significantly reduced ACE2-mediated virus infection. The S protein of SARS-CoV-1 that also uses human ACE2 also contains a positive residue (K) at this position, while the S proteins of CoVs utilizing other receptors vary at this location. Our results indicate that the presence of a positively charged amino acid at position 403 in the S protein is critical for efficient utilization of human ACE2. This finding could help to predict the zoonotic potential of animal coronaviruses.

---

Since its first occurrence in Wuhan in December 2019, SARS-CoV-2, the causative agent of COVID-19, has infected about 170 million people by May 2021 and caused a global health and economic crisis^3^. SARS-CoV-2 belongs to the *Sarbecovirus* subgenus of betacoronaviruses, which are mainly found in bats^3^. Horseshoe bats *(Rhinolophidae)* also harbour viruses that are closely related to SARS-CoV-1 that infected about 8.000 people in 2002 and 2003^3,4^. The bat virus RaTG13 sampled from a *Rhinolophus affinis* horseshoe bat in 2013 in Yunnan has been identified as the closest relative of SARS-CoV-2 showing approximately 96% sequence identity throughout the genome^1^. Thus, SARS-CoV-2 most likely originated from horseshoe bats^1,5^, although it has been proposed that cross-species transmission to humans may have involved pangolins as secondary intermediate host^6,7^.

The Spike (S) proteins of both SARS-CoV-1 and SARS-CoV-2 utilize the human angiotensin-converting enzyme 2 (ACE2) receptor to enter human target cells^8–11^. The ability to use a human receptor for efficient infection is a key prerequisite for successful zoonotic transmission. Although the RaTG13 S protein is highly similar to the SARS-CoV-2 S it does not interact efficiently with the human ACE2 receptor^2^, suggesting that this bat virus would most likely not be able to directly infect humans. It has been reported that specific alterations in the receptor-binding domain (RBD)^12^, as well as a four-amino-acid insertion (PRRA) introducing a furin-cleavage site,^8,13^ play a key role in efficient ACE2 utilization and consequently the high infectiousness and efficient spread of SARS-CoV-2. However, it remains poorly understood which specific features allow the S proteins of bat CoVs to use human ACE2 as entry cofactor and thus to successfully cross the species barrier to humans.

Computational analyses suggested that R403 is involved in intramolecular interactions stabilizing the SARS-CoV-2 S trimer interface^2^ and contributes significantly to the strength of SARS-CoV-2 RBD interaction with the human ACE2 receptor^14^. We found that R403 is highly conserved in SARS-CoV-2 S proteins: only 233 of 1.7 million sequence records contain a conservative change of R403K and just 18 another residue or deletion. Notably, the presence of a positively charged residue at position 403 distinguishes the S proteins of SARS-CoV-1 (K403) and SARS-CoV-2 (R403) from the bat CoV RaTG13 S protein (T403) (**Fig. 1a**). Molecular modelling of S/ACE2 interaction using reactive force field simulations confirmed close proximity and putative charge interactions between R403 in the SARS-CoV-2 S with E37 in the human ACE2 receptor (**Fig. 1b**). These analyses predicted that mutation of T403R significantly strengthens the ability of the RaTG13 S protein to bind human ACE2 (**Fig. 1c, Extended Data Movies 1 and 2**).

**Fig. 1:**
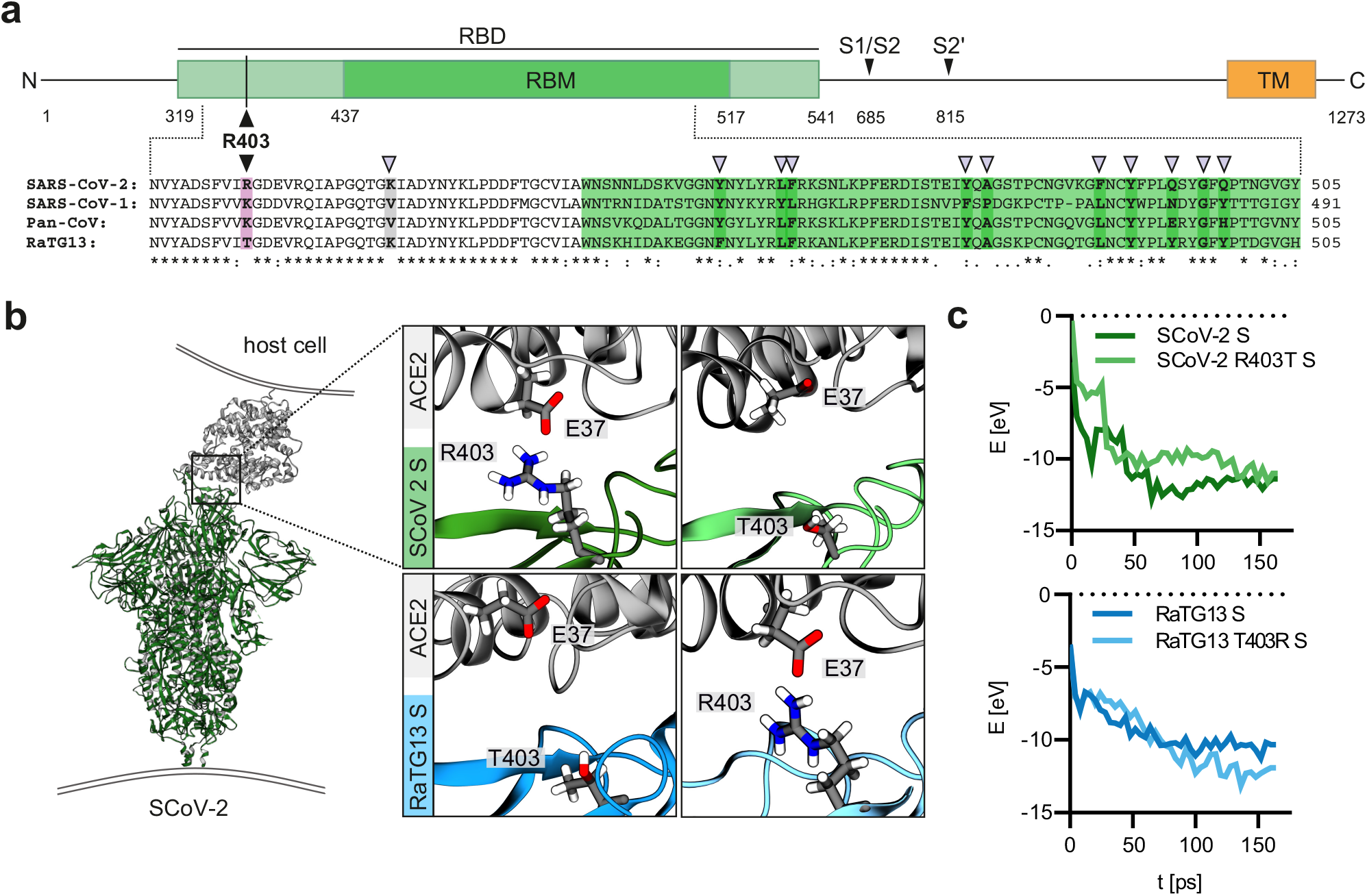
Modelling of the interaction of Coronavirus Spike residue 403 with human ACE2. **a,** Schematic representation of the SARS-CoV-2 S protein (top panel), domains are indicated in different colors. Receptor binding domain (RBD), light green. Receptor binding motif (RBM), dark green. Transmembrane domain (TM), orange. R403, pink. S1/S2 and S2’ cleavage sites are indicated. Sequence alignment of SARS-CoV-2, SARS-CoV-1, Pan-CoV and RaTG13 Spike RBD (bottom panel). Sequence conservation is indicated. purple arrows denote important residues for ACE2 binding. **b,** Reactive force field simulation of SARS-CoV-2 Spike in complex with human ACE2 (PDB: 7KNB) (left panel) and focus on position 403 in SARS-CoV-2 S (R) or RaTG13 S (T) or respective exchange mutants at position 403 (right panel). **c**, Exemplary energy curve of the reactive molecular dynamics simulation for SARS-CoV-2 S and SARS-CoV-2 S R403T (top panel) and RaTG13 and RaTG13 T430R spike with human ACE2 (bottom panel).

To verify the functional importance of residue 403 for ACE2 usage by CoV S proteins, we used VSV particles (VSVpp) pseudotyped with parental and mutant S proteins. Mutation of R403T reduced the ability of the SARS-CoV-2 S protein to mediate entry of VSVpp into Caco-2 cells by 40% (**Fig. 2a, b**). Strikingly, the T403R change enhanced the infectiousness of VSVpp carrying the RaTG13 S ~30-fold, while substitution of T403A introduced as control had no enhancing effect (**Fig. 2b**). Cell-to-cell fusion assays showed that coexpression of the SARS-CoV-2 S and human ACE2 resulted in the formation of large syncytia (**Extended Data Fig. 1**). The parental and T403A RaTG13 S did not lead to significant fusion but significant syncytia formation was observed for the T403R RaTG13 S (**Extended Data Fig. 1**). Western blot analyses showed that the mutant S proteins were efficiently expressed and incorporated into VSVpp, albeit the SARS-CoV-2 R403T S with reduced efficiency (**Extended Data Fig. 2**). In line with the VSVpp results, complementation of a full-length recombinant SARS-CoV-2 lacking the S ORF (SCoV-2ΔS) in ACE2-expressing HEK293T cells with wildtype (WT) SARS-CoV-2 S led to virus-induced cytopathic effects (CPE) indicating successful virus production and propagation (**Fig. 2c**). Mutation of R403T in the SARS-CoV-2 S reduced CPE. The WT and T403A RaTG13 S were entirely unable to complement SCoV-2ΔS, while the T403R RaTG13 S resulted in significant CPE. Expression of a Gaussia luciferase (GLuc) from S variant complemented recombinant SCoV2ΔS-GLuc confirmed the importance of R403 for viral spread (**Fig. 2d**).

**Fig. 2:**
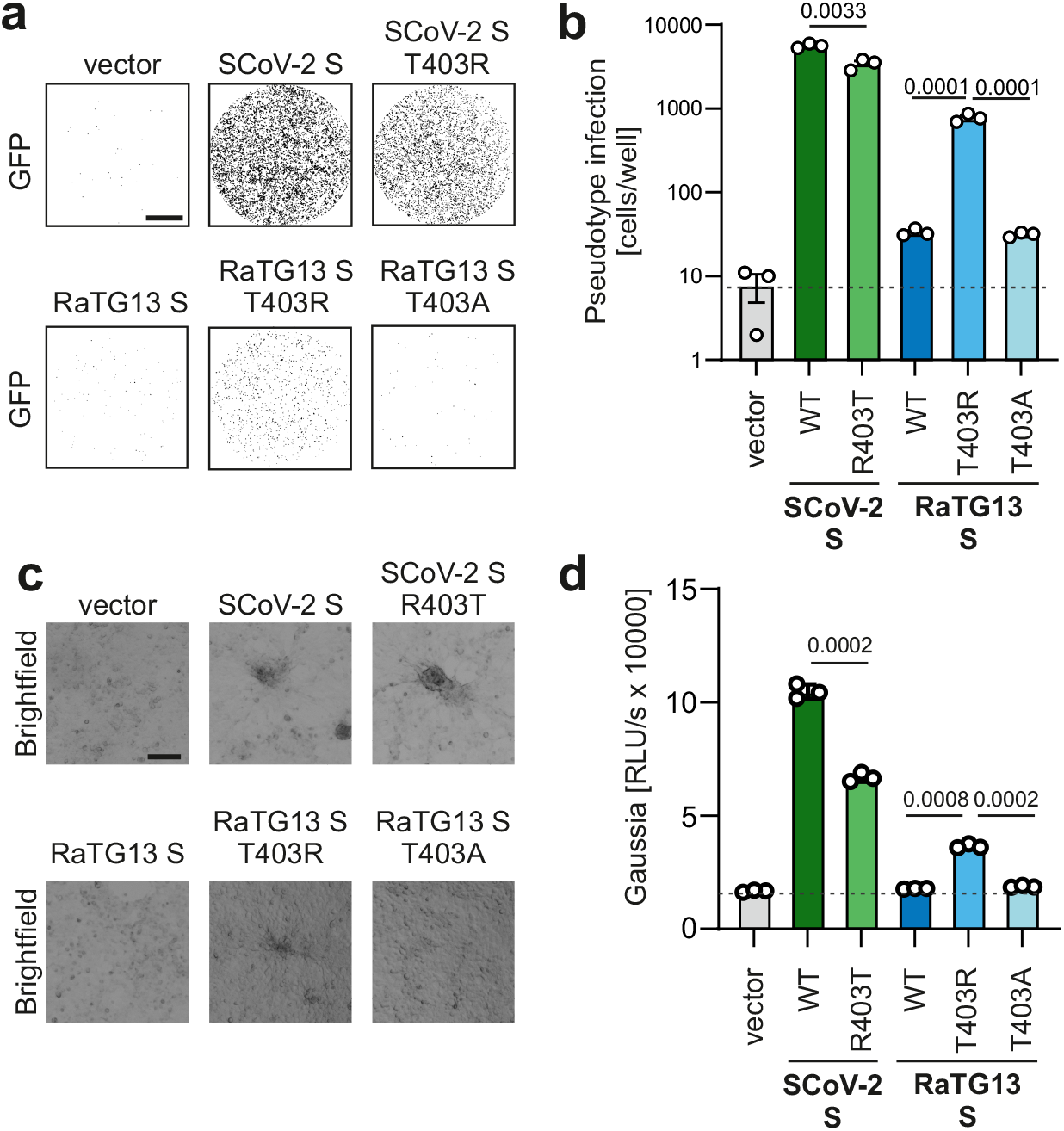
R403 in Spike is crucial to use ACE2 as an entry receptor. **a,** Binary images of CaCo2 cells transduced with VSVΔG-GFP pseudotyped with SARS-CoV-2, RaTG13 or indicated mutant S. Successful infection events (=GFP positive cells) displayed as black dots. Scale bar, 1.5mm. **b,** Automatic quantification of infection events by counting GFP positive cells. n=3 (biological replicates) ± SEM. **c,** Bright field and fluorescence microscopy (GFP) images of HEK293T cells transfected with SCoV-2ΔS bacmid, SCoV2-N, ACE2, T7 polymerase and indicated Spike variants. Scale bar, 125μm. **d,** Quantification of Gaussia luciferase activity in the supernatant of HEK293T cells expressing SCoV-2ΔS-Gaussia bacmids as described in (c). n=3 (biological replicates) ± SEM. P values are indicated (student’s t test).

Coronavirus entry is a multi-step process and critically dependent on proteolytic processing of the S protein^15^. The interaction of the SARS-CoV-2 S trimer with ACE2 promotes proteolytic processing^16,17^. Western blot analysis revealed that ACE2 coexpression induces efficient cleavage of the SARS-CoV-2 and T403R RaTG13 S proteins to S2, while cleavage of the WT and T403A RaTG13 S proteins remained inefficient (**Extended Data Fig. 3**). R403 generates a potential RGD integrin binding site in the viral Spike protein and it is under debate whether the ability of the SARS-CoV-2 S to use integrins as viral attachment factors may play a role in its high infectiousness^18,19^. The integrin inhibitor ATN-161 had no significant effect on SARS-CoV-2 or T403R RaTG13 S-mediated infection (**Extended Data Fig. 4a, b**). Thus, the enhancing effect of the T403R mutation on the ability of RaTG13 S to infect human cells seems to be due to increased interaction with ACE2 rather than utilization of integrins. Taken together, our results demonstrate that mutation of T403R strongly enhances the ability of the bat RaTG13 S protein to utilize ACE2 for infection of human cells.

To assess whether the T403R change might allow the bat CoV RaTG13 to spread to different human organs, we performed infection studies using intestinal organoids derived from pluripotent stem cells. The parental SARS-CoV-2 S protein allowed efficient infection of gut organoids^20^ and the R403T change had modest attenuating effects (**Fig. 3, Extended Data Fig. 5**). In contrast, the parental RaTG13 S protein did not result in significant VSVpp infection, while the corresponding T403R mutant allowed significant infection of human intestinal cells (**Fig. 3; Extended Data Fig. 5**).

**Fig. 3:**
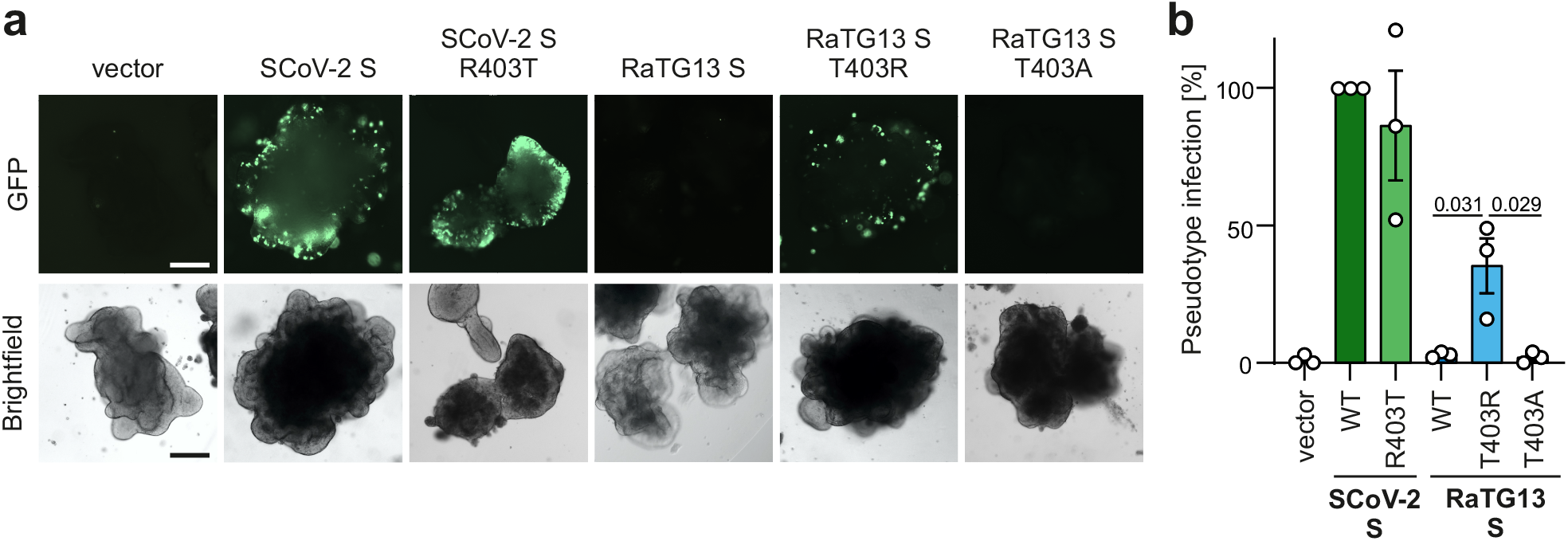
T403R allows RaTG13 S to mediate infection of human gut organoids. **a,** Bright field and fluorescence microscopy (GFP) images of hPSC derived gut organoids infected with VSVΔG-GFP (green) pseudotyped with SARS-CoV-2, RaTG13 or indicated mutant S (300 μl, 2 h). Scale bar, 250μm. **b,** Quantification of the percentage of GFP-positive cells of (a). n=3 (biological replicates) ± SEM. P values are indicated (student’s t test).

To examine the species-specificity of receptor usage by SARS-CoV-2 and RaTG13 S proteins, we overexpressed human and bat derived ACE2 in HEK293T cells and examined their susceptibility to S-mediated VSVpp infection. The WT SARS-CoV-2 and the T403R RaTG13 S proteins allowed efficient entry into cells overexpressing human ACE2, while the parental RaTG13 S protein was poorly active (**Fig. 4a**). Both WT SARS-CoV-2 S and (to a lesser extent) R403T SARS-CoV-2 S proteins were also capable of using bat *(Rhinolophus affinis)* ACE2 for viral entry although the overall infection rates were low (**Fig. 4a, Extended Data Fig. 6**). In contrast, the RaTG13 S proteins were unable to use bat ACE2 for infection suggesting that RaTG13 might use an alternative receptor for infection of bat cells. The results agree with the previous finding that RaTG13 S is able to use human ACE2 to some extent if overexpressed^21^ but further demonstrate that the T403R greatly enhances this function and is required for utilization of endogenously expressed human ACE2.

**Fig. 4:**
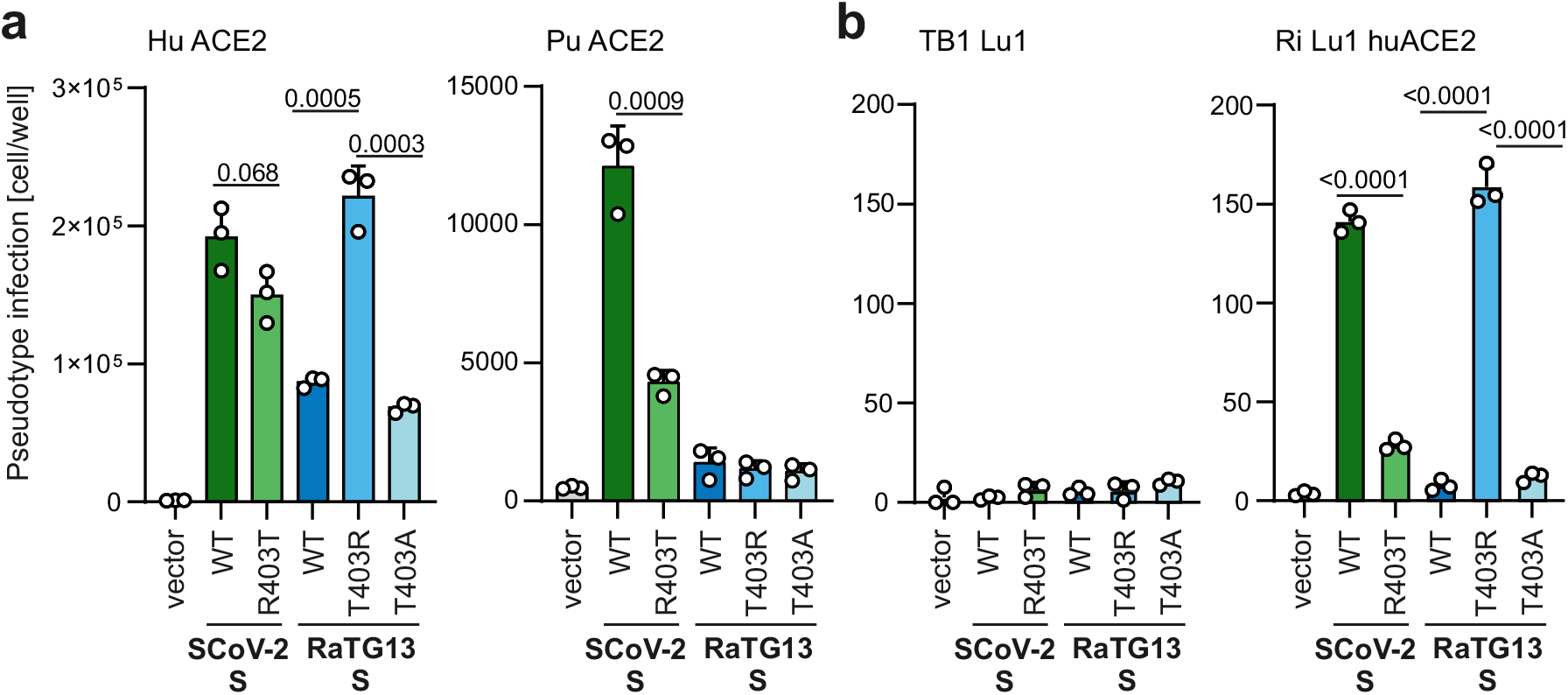
SARS-CoV-2 S and T403R RaTG13 S allow entry with human but not bat ACE2. **a,** HEK293T cells expressing indicated ACE2 (Human ACE2 or *Rhinolophus affinis* ACE2) constructs or **b,** Tb 1 Lu, *Tadarida brasiliensis* derived lung epithelial and Ri 1 Lu huACE2 *Rhinolophus affinis* derived lung epithelial cells expressing human ACE2 were infected with VSVΔG-GFP pseudotyped with SARS-CoV-2, RaTG13 or indicated mutant S. Quantification by automatic counting of GFP positive cells. n=3 (biological replicates) ± SEM. P values are indicated (student’s t test).

To validate the results obtained with human HEK293T cells, we utilized the lung epithelial cell line Tb1 Lu1 of *Tadarida brasiliensis* (Bat31)^22^. In agreement with the previous finding that this cell line lacks endogenous ACE2 expression, it did not support infection by CoV S proteins (**Fig. 4b**). Engineered expression of human ACE2 rendered Lu 1 highly susceptible to infection mediated by SARS-CoV-2 and the T403R RaTG13 S proteins (**Fig. 4b**). In comparison, entry via the R403T SARS-CoV-2 S was strongly attenuated and the parental and T403A RaTG13 S proteins were unable to mediate significant VSV-pp infection.

Our results demonstrate that a single amino acid change of T403R allows RaTG13, the closest known bat relative of SARS-CoV-2, to utilize human ACE2 for viral entry. The strong enhancing effect of the T403R change on RaTG13 S function came as surprise since five of six different residues proposed to be critical for SARS-CoV-2 S RBD interaction with human ACE2 are not conserved in RaTG13 S^12,23^. A very recent study proposed that residue 501 plays a key role in the ability of RaTG13 S to use human ACE2 for viral entry^24^ but the reported enhancing effect of changes at position 501 was weaker than that observed for the T403R change analysed in the present study. However, the previous finding that numerous residues in the SARS-CoV-2 S RBD are involved in the interaction with the human ACE2 orthologue explains why the R403T substitution only moderately reduced SARS-CoV-2 infection. It has been shown that the RBD of SARS-CoV-2 S shows higher homology to the corresponding region of the pangolin CoV S protein than to RaTG13^6,7^. Whether or not this is a consequence of recombination or convergent evolution is under debate^25,26^. Notably, the Pan CoV-S protein also contains a positive residue (K) at position 403 (**Fig. 1a**) and is capable of utilizing human ACE2 for infection. Altogether our results suggest that a positive residue at position 403 in the S protein was most likely a prerequisite for efficient zoonotic transmission and pandemic spread of SARS-CoV-2. We found that a positively charged residue at the corresponding position is present in the S proteins of the great majority of RaTG13-related bat coronaviruses (**Extended Data Fig. 7**) raising the possibility that many bat sarbecoviruses, including the unknown precursor of SARC-CoV-2, are fitter for zoonotic transmission than RaTG13.

## Methods

### Molecular dynamics simulation

Based on the structure of ACE2-bounded to SARS-CoV-2 taken from the Protein Data Bank^27^ (identification code 7KNB), the initial atomic positions were obtained. Equilibration (300K for 0.5 ns) was performed by ReaxFF (reactive molecular dynamic) simulations^28^ within the Amsterdam Modeling Suite 2020 (ADF2020, SCM, Theoretical Chemistry, Vrije Universiteit, Amsterdam, The Netherlands, http://www.scm.com). Based on the equilibrated structure, amino acids from the spike protein were replaced with the respective amino acids from RaTG13, respectively the modification. After an additional equilibration (300K for 0.5 ns) ReaxFF (reactive molecular dynamic) simulations were performed within the *NVT* ensemble over 25 ps, while coupling the system to a Berendsen heat bath (T=300 K with a coupling constant of 100 fs). The interaction energy was obtained by averaging over these simulations. For all visualizations the Visual Molecular Dynamics program (VMD)^29^ was used.

### Cell culture and viruses

All cells were cultured at 37°C in a 5% CO2 atmosphere. Human embryonic kidney 293T cells purchased from American type culture collection (ATCC: #CRL3216) were cultivated in Dulbecco’s Modified Eagle Medium (DMEM, Gibco) supplemented with 10% (v/v) heat-inactivated fetal bovine serum (FBS, Gibco), 2 mM L-glutamine (PANBiotech), 100 μg/ml streptomycin (PANBiotech) and 100 U/ml penicillin (PANBiotech). Calu-3 (human epithelial lung adenocarcinoma, kindly provided and verified by Prof. Frick, Ulm University) cells were cultured in Minimum Essential Medium Eagle (MEM, Sigma) supplemented with 10% (v/v) FBS (Gibco) (during viral infection) or 20% (v/v) FBS (Gibco) (during all other times), 100 U/ml penicillin (PAN-Biotech), 100 μg/ml streptomycin (PAN-Biotech), 1 mM sodium pyruvate (Gibco), and 1 mM NEAA (Gibco). Caco-2 (human epithelial colorectal adenocarcinoma, kindly provided by Prof. Holger Barth, Ulm University) cells were cultivated in DMEM (Gibco) containing 10% FBS (Gibco), 2 mM glutamine (PANBiotech), 100 μg/ml streptomycin (PANBiotech), 100 U/ml penicillin (ANBiotech), 1 mM Non-essential amino acids (NEAA, Gibco), 1 mM sodium pyruvate (Gibco). I1-Hybridoma cells were purchased from ATCC (#CRL-2700) and cultured in RPMI supplemented with 10% (v/v) heat-inactivated FBS (Gibco), 2 mM L-glutamine (PANBiotech), 100 μg/ml streptomycin (PANBiotech) and 100 U/ml penicillin (PANBiotech). Tb 1 Lu *(Tadarida brasiliensis* derived lung epithelial) and Ri 1 Lu huACE2 *(Rhinolophus affinis* derived lung epithelial cells expressing human ACE2, ACE2, kindly provided by Marcel A. Müller, were cultured in DMEM supplemented with 10% (v/v) heat-inactivated FBS (Gibco), 2 mM L-glutamine (PANBiotech), 100 μg/ml streptomycin (PANBiotech) and 100 U/ml penicillin (PANBiotech), 2 mM sodium pyruvate (Gibco). Viral isolate BetaCoV/France/IDF0372/2020 (#014V-03890) was obtained through the European Virus Archive global.

### Expression constructs

pCG_SARS-CoV-2-Spike-IRES_eGFP, coding the spike protein of SARS-CoV-2 isolate Wuhan-Hu-1, NCBI reference Sequence YP_009724390.1, was kindly provided by Stefan Pöhlmann (German Primate Center, 473 Göttingen, Germany). pCG_SARS-CoV-2-Spike C-V5-IRES_eGFP and RaTG13-S (synthesized by Baseclear) was PCR amplified and subcloned into a pCG-IRES_eGFP expression construct using the restriction enzymes XbaI and MluI (New England Biolabs). The SARS-CoV-2 S R403T and RaTG13 S T403R/T403A mutant plasmids were generated using Q5 Site-Directed Mutagenesis Kit (NEB).

### Cloning of SARS-CoV-2 ΔS bacmid

An anonymized residual respiratory swab sample from a patient with SARS-CoV-2 infection was used as a template for genome amplification. Total nucleic acids were extracted on an automated Qiagen EZ1 robotic workstation using the Qiagen EZ1 virus mini kit v2.0 according to the manufacturer’s instructions. Genomic viral RNA was reverse transcribed using the NEB LunaScript RT SuperMix Kit according to the manufacturer’s protocol. Four overlapping fragments covering the entire viral genome were amplified using the NEB Q5 High-Fidelity DNA Polymerase. The resulting amplicons were assembled with a modified pBeloBAC11 backbone, containing CMV and T7 promotors as well as the HDV ribozyme and bGH polyA signal, using the NEBuilder HiFi DNA Assembly Cloning Kit. Assembled DNA was electroporated into *E. coli* GS1783 strain and resulting clones of pBelo-SARSARS-CoV-2 were confirmed by restriction digestion and next generation sequencing. The viral Spike gene was replaced with a kanamycin-cassette flanked by SacII restriction sites by homologous recombination using the Lambda-Red Recombination System^30^. The bacmid was linearized with the restriction enzyme SacII, and EGFP or GLuc reporter cassettes were introduced instead of Spike using the the NEBuilder HiFi DNA Assembly Cloning Kit according to the manufacturer’s instruction. Positive clones were confirmed by restriction digestion and sequencing.

### SARS-CoV-2 ΔS replicon system

HEK293 T cells were seeded in six well format and transfected with 3 μg pBelo-SARSCoV-2-dSpike-GLuc-K2 or pBelo-SARSCoV-2-dSpike-EGFP and 0.25 μg of each expression construct pLVX-EF1alpha-SARS-CoV2-N-2xStrep-IRES-Puro, pCG-ACE2, pCAG-T7-RNA-polymerase and one pCG-vector encoding the spike protein of SARS-CoV-2, RaTG13 or the indicated mutant S respectively. Two days after transfection, bright field and fluorescence microscopy (GFP) images were acquired using the Cytation 3 microplate reader (BioTek). Gaussia luciferase activity in the supernatants was measured with the Gaussia Luciverase Assay system (Promega) according to the company’s instructions.

### Transfections

Plasmid DNA was transfected using either calcium phosphate transfection or Polyethylenimine (PEI, 1 mg/ml in H2O, Sigma-Aldrich) according to the manufacturers recommendations or as described previously^31^.

### Pseudoparticle production

To produce pseudotyped VSVΔG-GFP particles, 6*10^6^ HEK 293 T cells were seeded 18 hours before transfection in 10 cm dishes. The cells were transfected with 15 μg of a glycoprotein expressing vector using PEI (PEI, 1 mg/ml in H2O, Sigma-Aldrich). Twenty-four hours post transfection, the cells were infected with VSVΔG-GFP particles pseudotyped with VSV G at a MOI of 3. One hour post-infection, the inoculum was removed. Pseudotyped VSVΔG-GFP particles were harvested 16 hours post infection. Cell debris were pelleted and removed by centrifugation (500 g, 4 °C, 5 min). Residual input particles carrying VSV-G were blocked by adding 10 % (v/v) of I1 Hybridoma Supernatant (I1, mouse hybridoma supernatant from CRL-2700; ATCC) to the cell culture supernatant.

### Whole-cell and cell free lysates

Whole-cell lysates were prepared by collecting cells in Phosphate-Buffered Saline (PBS, Gibco), pelleting (500 g, 4 °C, 5 min), lysing and clearing as previously described^31^. Total protein concentration of the cleared lysates was measured using the Pierce BCA Protein Assay Kit (Thermo Scientific) according to manufacturer’s instructions. Viral particles were filtered though a 0.45 μm MF-Millipore Filter (Millex) and centrifuged through a 20% sucrose (Sigma) cushion. The pellet was lysed in transmembrane lysis buffer already substituted with Protein Sample Loading Buffer (LI-COR).

### SDS-PAGE and immunoblotting

SDS-PAGE and immunoblotting was performed as previously described^31^. In brief, whole cell lysates were mixed with 4x Protein Sample Loading Buffer (LI-COR, at a final dilution of 1x) supplemented with 10% (v/v) Tris(2-Carboxyethyl)phosphine hydrochloride 0.5 M (SIGMA), heated to 95°C for 10 min separated on NuPAGE 4-12% Bis-Tris Gels (Invitrogen) for 90 min at 120 V and blotted at constant 30 V for 30 min onto Immobilon-FL PVDF membrane (Merck Millipore). After the transfer, the membrane was blocked in 1% Casein in PBS (Thermo Scientific) and stained using primary antibodies directed against SARS-CoV-2 S (1:1,000, Biozol, 1A9, #GTX632604), ACE2 (1:1,000, Abcam, #GTX632604), VSV-M (1:2,000, Absolute Antibody, 23H12, #Ab01404-2.0), V5-tag (1:1,000, Cell Signaling, #13202), GAPDH (1:1,000, BioLegend, #631401) and Infrared Dye labelled secondary antibodies (1:20,000, LI-CORIRDye). Proteins were detected using a LI-COR Odyssey scanner and band intensities were quantified using LI-COR Image Studio.

### Stem Cell Culture and Intestinal Differentiation

Human embryonic stem cell line HUES8 (Harvard University, Cambridge, MA) was used with permission from the Robert Koch Institute according to the “79. Genehmigung nach dem Stammzellgesetz, AZ 3.04.02/0084.” Cells were cultured on human embryonic stem cell matrigel (Corning, Corning, NY) in mTeSR Plus medium (STEMCELL Technologies, Vancouver, Canada) at 5% CO2, 5% O2, and 37°C. Medium was changed every other day and cells were split with TrypLE Express (Invitrogen, Carlsbad, CA) twice a week. For differentiation, 300,000 cells per well were seeded in 24-well plates coated with growth factor–reduced matrigel (Corning) in mTeSR Plus with 10 mM Y-27632 (STEMCELL Technologies). The next day, differentiation was started at 80%-90% confluency, as described previously^32^.

### Intestinal organoids

To prepare in vitro differentiated organoids for transduction, matrigel was dissolved in Collagenase/Dispase (Roche, Basel, Switzerland) for 2 hours at 37°C and stopped by cold neutralization solution (DMEM, 1% bovine serum albumin, and 1% penicillinstreptomycin). Organoids were transferred into 1.5-mL tubes and infected in 300 μL pseudoparticle containing inoculum. Organoids were then resuspended in 35-μL cold growth factor-reduced matrigel to generate cell-matrigel domes in 48-well plates. After 10 minutes at 37°C, intestinal growth medium (DMEM F12 [Gibco, Gaithersburg, MD], 1× B27 supplement [Thermo Fisher Scientific], 2-mM L-glutamine, 1% penicillin-streptomycin, 40 mM HEPES [Sigma-Aldrich], 3 μM CHIR99021, 200 nM LDN-193189 [Sigma-Aldrich], 100 ng/mL hEGF [Novoprotein, Summit, NJ], and 10 μM Y-27632 [STEMCELL Technologies]) was added and organoids were incubated at 37°C. The Organoids were imaged using the Cytation 3 cell imaging system and processed with Gen 5 and ImageJ software. For FACS preparation, the matrigel was dissolved and the extracted organoids were dissolved in Accutase (Stemcell technologies). The cells were fixed with PBS for 10 min at 4°C and washed with cold PBS containing 2% FBS. Flow cytometry analyses were performed using a FACS CANTO II (BD) flow cytometer. Transduction rates were determined by GFP expression and analysed with DIVA and Flowjow10 software.

### α5β5 integrin blocking

Caco-2 cells were preincubated with the indicated amounts of α5β5 integrin Inhibitor ATN-161 (Sigma) for two hours and infected with 100 μl freshly produced VSVΔG-GFP pseudo particles. 16 hours post infection, GFP positive cells were automatically quantification using a Cytation 3 microplate reader (BioTek). Calu-3 cells were preincubated with the indicated amounts of ATN-161 (Sigma) for two hours and infected with SARS-CoV-2 Viral isolate BetaCoV/France/IDF0372/2020 (MOI 0.05, six hours). 48 hours post-infection supernatants were harvested for qRT-PCR analysis.

### Sequence Logo and alignments

Alignments of primary bat sequences (GQ153541.1/1-71,GQ153544.1/1-71,GQ153540.1, GQ153539.1, DQ084200.1, DQ084199.1, GQ153548.1, GQ153547.1, GQ153546.1, GQ153545.1, DQ022305.2, GQ153542.1, GQ153543.1, KJ473815.1, KF294457.1, KY417148.1, KJ473814.1, MK211374.1, KY417142.1, MK211377.1, JX993988.1, DQ412043.1, DQ648857.1, JX993987.1, KY417143.1, KY417147.1, MK211378.1, DQ648856.1, KJ473812.1, KY770860.1, KY770858.1, KY770859.1, KJ473816.1, RmYN02, KY417145.1, KU182964.1, KY938558.1, KJ473811.1, KJ473813.1, MG772933.1, MG772934.1, KY417150.1, KT444582.1, KY417152.1, MK211376.1, GU190215.1, MN996532.1, EF065513.1, MG693170.1, MG762674.1, HM211101.1, HM211099.1, EF065514.1, EF065516.1, EF065515.1, MK492263.1, MG693168.1, MG693172.1, MG693169.1, MG693171.1, KU762337.1, KU762338.1, HQ166910.1, KT253270.1, KT253269.1, KY073748.1, MN611517.1, KY073747.1, KY073744.1, KY073745.1, KY073746.1, NC_028833.1, MK720944.1, NC_010437.1/1-7,EU420138.1, KJ473796.1, MN611524.1, KJ473795.1, EU420137.1, KJ473799.1, KJ473800.1, KJ473797.1, MN611518.1, KY770850.1, KY770851.1, KJ473806.1, EU420139.1, KJ473798.1, MG916902.1, MG916903.1, JQ989269.1, JQ989267.1, JQ989268.1, JQ989266.1, JQ989272.1, JQ989273.1, MN611523.1, MN611525.1, JQ989271.1, JQ989270.1, MK720945.1, MK720946.1, MG916904.1, KJ473810.1, NC_028814.1, DQ648858.1, NC_009657.1, MN611521.1, KF430219.1, NC_009988.1/1-7,EF203066.1, EF203067.1, EF203065.1, MF370205.1, KJ473808.1, MN611522.1, DQ648794.1, EF065505.1, EF065506.1, EF065508.1, MH002339.1, MN611519.1, MH002338.1, KJ473822.1, MH002337.1, KU182965.1, EF065507.1, EF065510.1, EF065511.1, EF065512.1, MH002342.1, EF065509.1, KJ473820.1, MH002341.1, MN611520.1, KX442565.1, KX442564.1/1-71) was performed using ClustalW^33^ with a gapOpening penalty of 80. Sequence logos were generated using R packages ggplot2 and ggseqlogo^34^.

### Statistics

Statistical analyses were performed using GraphPad PRISM 8 (GraphPad Software). P-values were determined using a two-tailed Student’s t test with Welch’s correction. Unless otherwise stated, data are shown as the mean of at least three independent experiments ± SEM.

## Acknowledgments

We thank Kerstin Regensburger, Regina Burger, Jana-Romana Fischer, Birgit Ott, Martha Meyer, Nicole Schrott and Daniela Krnavek for technical assistance. The ACE2 vector and the SARS-CoV-2 S-HA plasmid were kindly provided by Shinji Makino and Stefan Pöhlmann, and bat cells by Marcel A. Müller. F.Z., C.P.B., J.K. and L.K. are part of the International Graduate school for Molecular Medicine (IGradU), Ulm. This study was supported by DFG grants to F.K. (CRC 1279, SPP 1923), K-K.C. (Co260/6-1 Neuro-COVID), T.J. (CRC1279), A.K. (KL 2544/8-1, KL 2544/5-1,7-1 and the ‘Heisenberg-Programm’ KL 2544/6-1) and K.M.J.S. (CRC1279, SP1600/6-1). A.E. is funded by the State of Bavaria “BAY-VoC” and “Coronaforschung”. F.K., K.M.J.S. and A.E. were supported by the BMBF (Restrict SARS-CoV-2, IMMUNOMOD and 01KI20172A SENSE-CoV2).

## Author Contributions

F.Z. performed most experiments. D.S., M.V., Q.X. and L.K. performed western blots and interaction assays. J.K., S.H. and A.K. generated and provided gut organoids. C.J. and T.J. performed molecular modelling analyses. K.-K.C. provided pseudotypes and reagents. F.Z., D.S., K.M.J.S. and F.K. conceived the study, planned experiments and wrote the manuscript. All authors reviewed and approved the manuscript.

## Materials & Correspondence

Further information and requests for resources and reagents should be directed to and will be fulfilled by Frank Kirchhoff.

## Extended Figure legends

**Extended Data Fig. 1:**
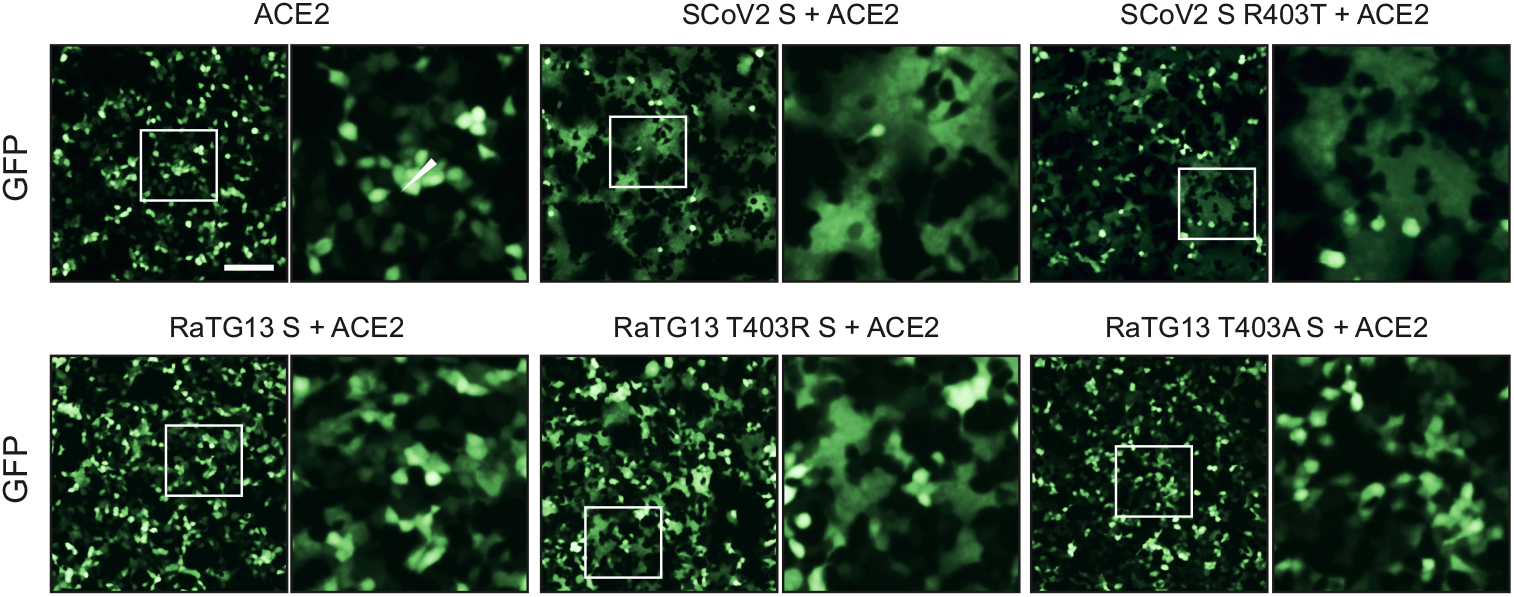
T403R RaTG13 S allows ACE2 dependent cell fusion. Exemplary fluorescence microscopy images of HEK293T cells expressing SCoV2 S, RaTG13 S or the indicated mutant, Human ACE2 and GFP (green). Insets are indicated by white boxes. Scale bar, 125μm.

**Extended Data Fig. 2:**
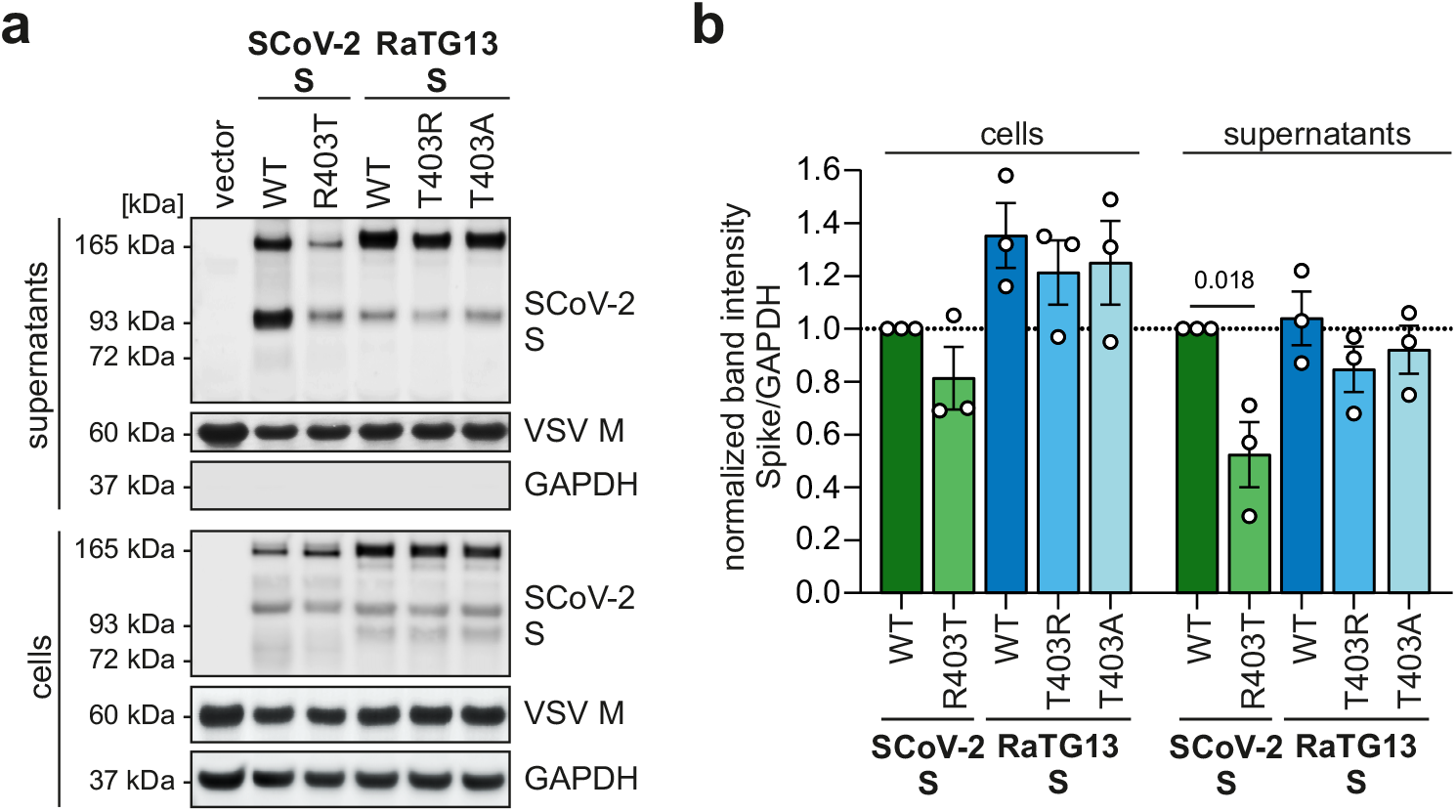
Incorporation of Spike variants in VSV pseudoparticles. **a,** Exemplary immunoblots of whole cells lysates (WCLs) and supernatants of HEK293T cells expressing SCoV2 S, RaTG13 S or the indicated mutant that were infected with VSVΔG-GFP. Blots were stained with anti-SARS-CoV-2 S, anti-GAPDH and anti-VSV-M. **b,** Quantification of Spike expression. n=3 (biological replicates) ± SEM. P values are indicated (student’s t test).

**Extended Data Fig. 3:**
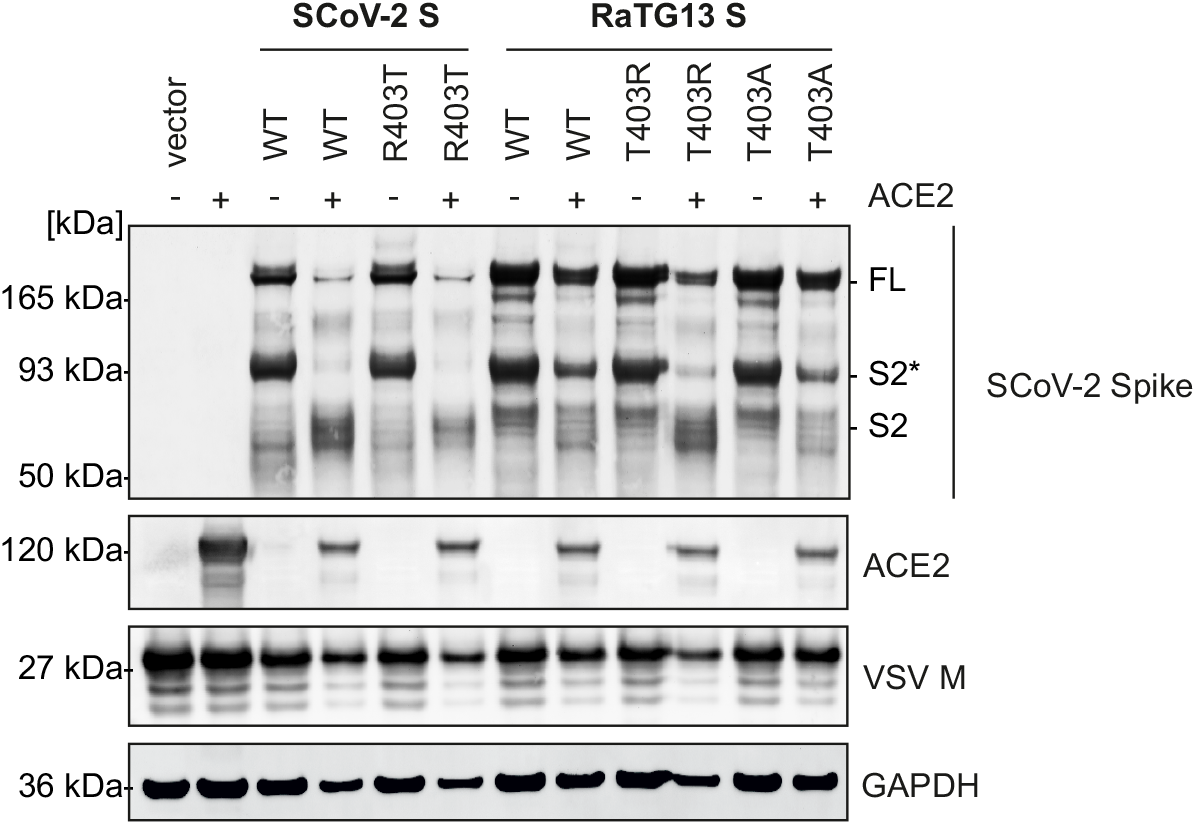
Processing of Spike proteins by ACE2 expression. **a,** Exemplary immunoblots of WCLs of HEK293T cells expressing SARS-CoV-2 S, RaTG13 S or the indicated mutant coexpressing Human ACE2 or empty vector construct. The blots were stained with anti-SARS-CoV-2 S, anti-GAPDH, anti-ACE2 and anti-VSV-M.

**Extended Data Fig. 4:**
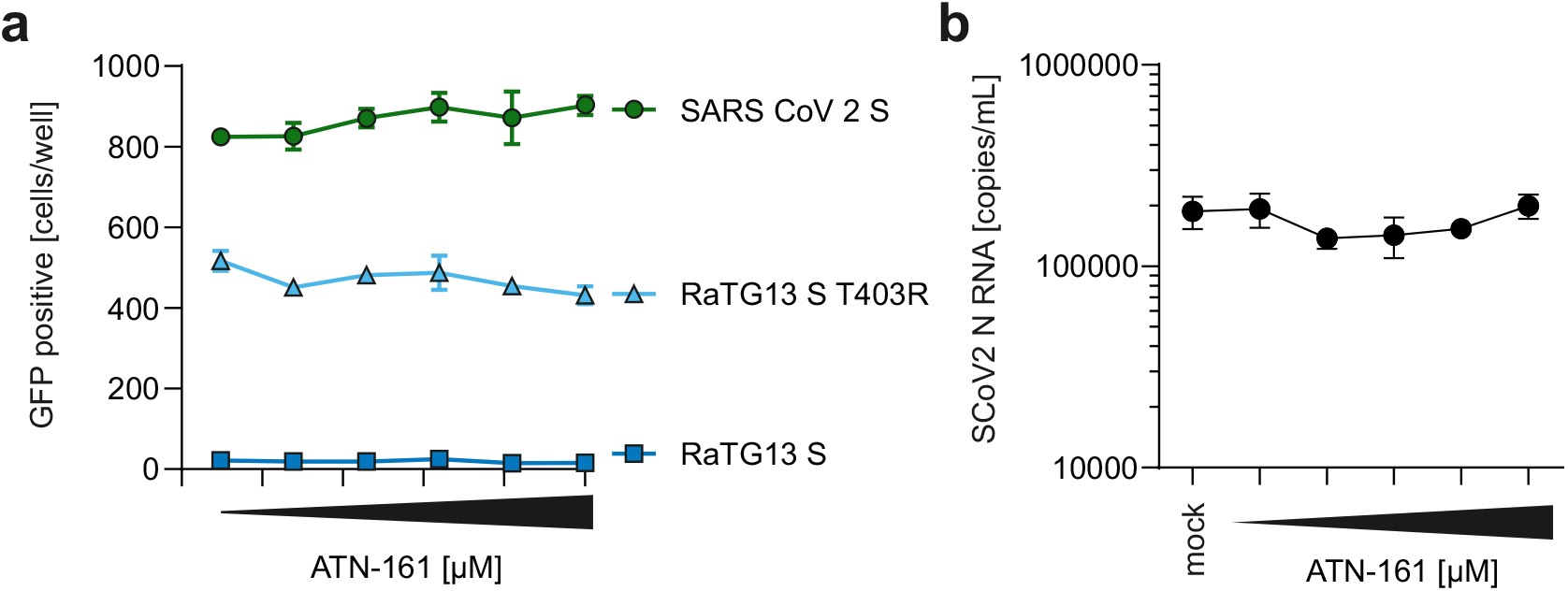
SARS-CoV-2 entry is independent of α5β5 integrin. **a,** Automated quantification by GFP fluorescence of Caco-2 cells preincubated with indicated amounts of α5β5 integrin Inhibitor ATN-161 and infected with VSVΔG-GFP pseudotyped with SARS-CoV-2, RaTG13 T403R mutant or RaTG13 S. n=3 (biological replicates) ± SEM. **b,** Quantification of viral RNA copies in the supernatant of Calu-3 cells preincubated with indicated amounts of ATN-161 and infected SARS-CoV-2 (MOI 0.05, 6 h). n=3 (biological replicates) ± SEM. P values are indicated (student’s t test).

**Extended Data Fig. 5:**
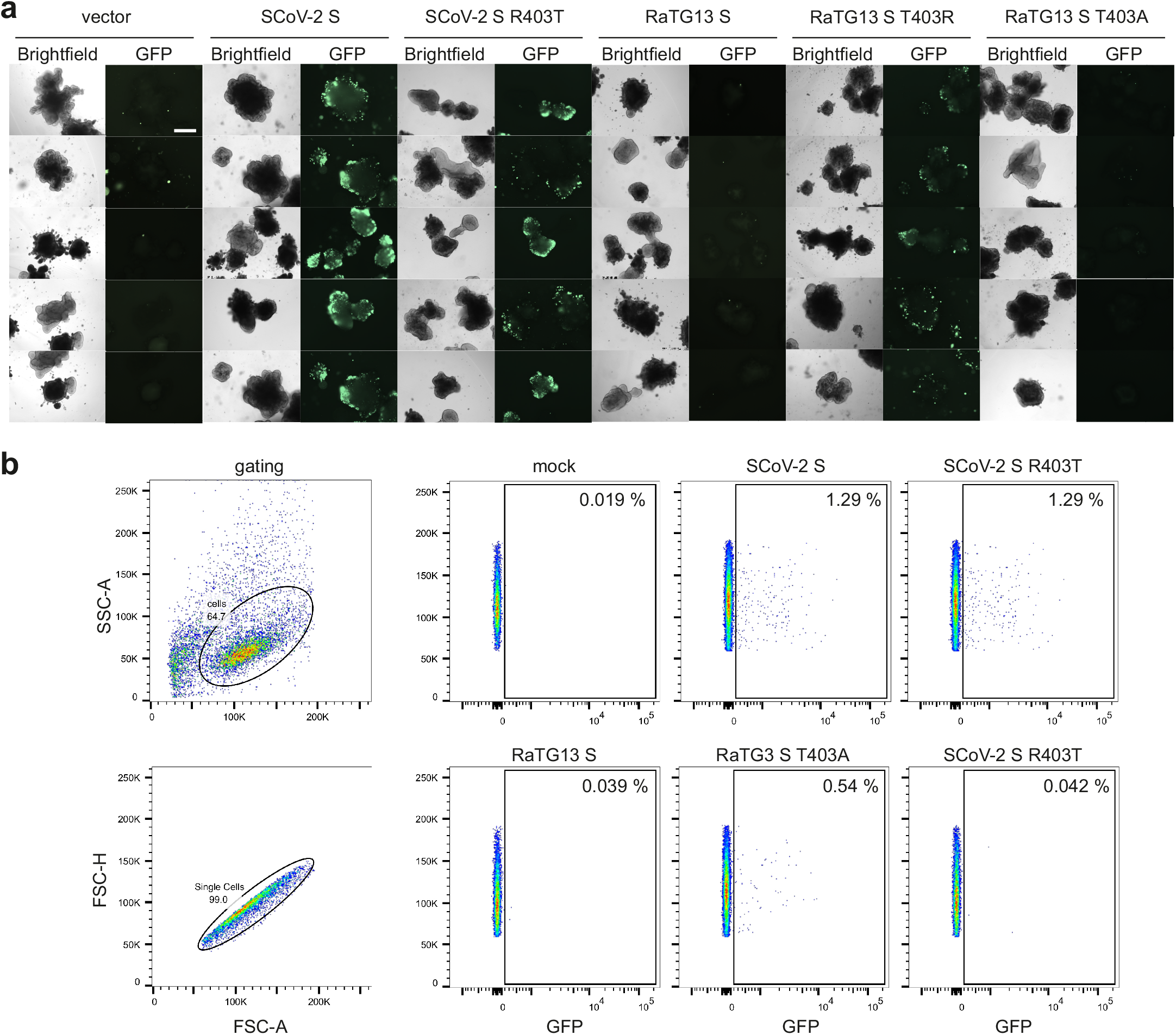
T403R allows RaTG13 S to mediate infection of human intestinal organoids. **a,** Bright field and fluorescence microscopy (GFP) images of hPSC derived gut organoids infected with equal amounts of VSVΔG-GFP (green) pseudotyped with SARS-CoV-2, RaTG13 or indicated mutant S (2 h). Scale bar, 250μm. **b,** Exemplary gating strategy of flow cytometry-based analysis of GFP-positive cells of (a). **c,** Quantification and exemplary gating. n=3 (biological replicates) ± SEM. P values are indicated (student’s t test).

**Extended Data Fig. 6:**
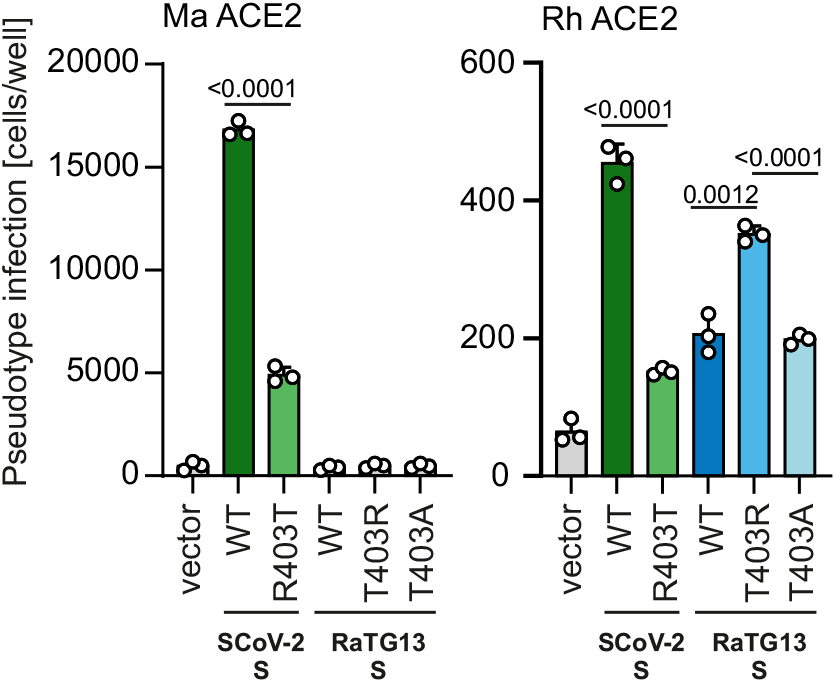
Bat ACE2 can be used for entry by SARS-CoV-2 Spike. Quantification of GFP positive HEK293T cells expressing indicated ACE2 variants *(Rhinolophus macrotis* ACE2 or *Rhinolophus rhodesiae* ACE2) infected with VSVΔG-GFP pseudotyped with SARS-CoV-2, RaTG13 or indicated mutant S. n=3 (biological replicates) ± SEM. P values are indicated (student’s t test).

**Extended Data Fig. 7:**
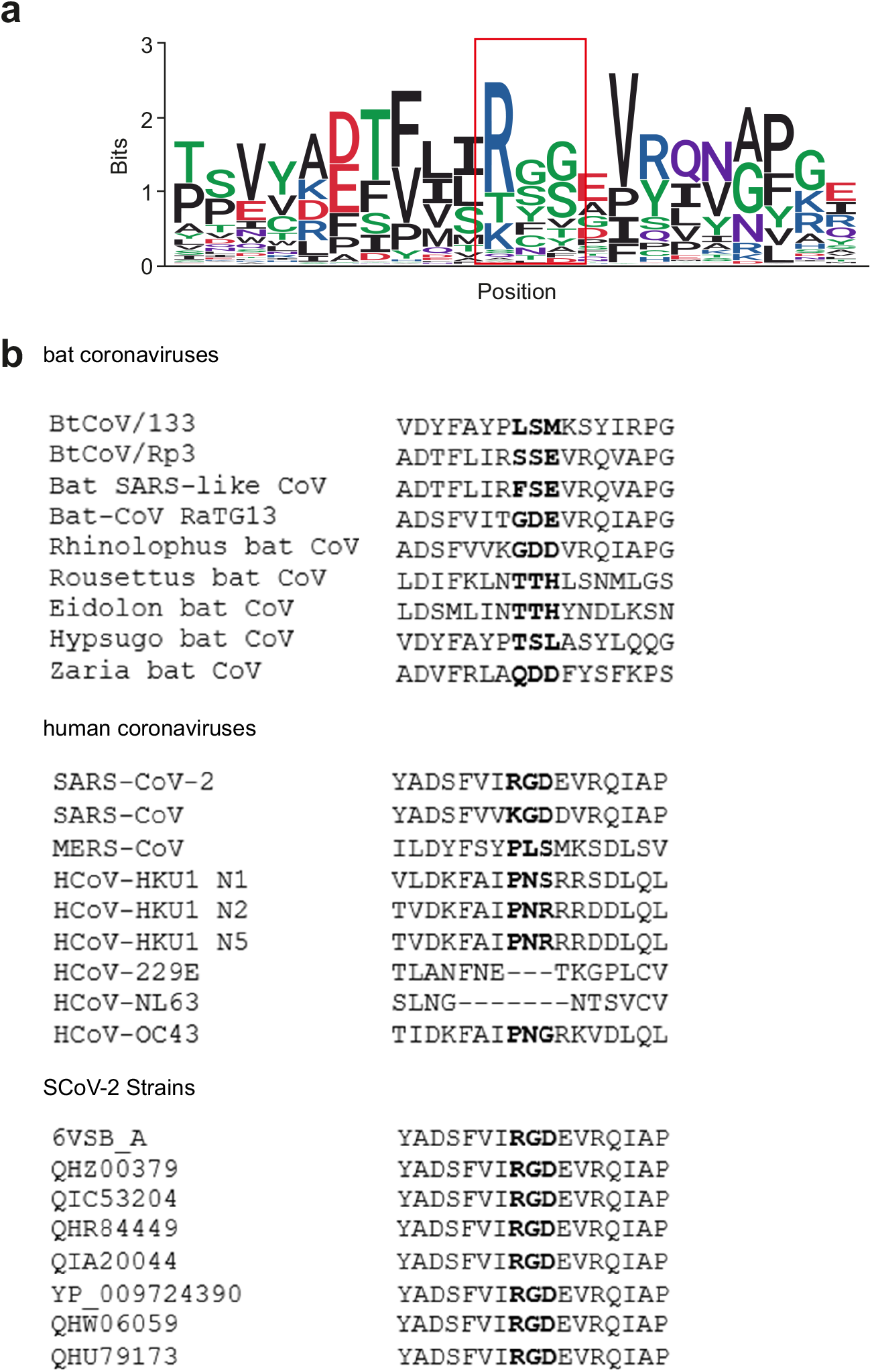
Conservation of the RGD motif in bat Coronavirus Spike proteins. **a,** Sequence logo of the alignment of 137 different bat Coronavirus Spike RBD sequences. The RGD motif is highlighted by a red box. **b,** Primary sequence alignment of selected bat coronaviruses, human coronaviruses and SARS-CoV-2 strains. The RGD motif is highlighted in bold.

